# Effects of the investigational drug sodium phenylbutyrate-TUDCA (AMX0035) on the transcriptional and metabolic landscape of sporadic ALS fibroblasts

**DOI:** 10.1101/2022.05.02.490306

**Authors:** Jasmine A. Fels, Jalia Dash, Kent Leslie, Giovanni Manfredi, Hibiki Kawamata

**Affiliations:** Feil Family Brain and Mind Research Institute, Weill Cornell Medicine, 407 East 61st Street, New York, NY 10065; Neuroscience Graduate Program, Weill Cornell Graduate School of Medical Sciences, 1300 York Ave, New York, NY 10065; Amylyx Pharmaceuticals, 43 Thorndike Street, Cambridge, Massachusetts; California Institute of Technology, Pasadena, CA 91106

**Keywords:** ALS, fibroblasts, AMX0035, Turso, Phenylbutyrate, WGCNA, RNAseq, metabolomics

## Abstract

ALS is a rapidly progressive, fatal disorder caused by motor neuron degeneration, for which there is a great unmet therapeutic need. AMX0035, a combination of sodium phenylbutyrate (PB) and taurursodiol (TUDCA, Turso), has shown promising results in early ALS clinical trials, but its mechanisms of action remain to be elucidated. To obtain an unbiased landscape of AMX0035 effects we investigated the transcriptomic and metabolomic profiles of primary skin fibroblasts from sporadic ALS patients and healthy controls treated with PB, TUDCA, or PB-TUDCA combination (Combo). Combo changed many more genes and metabolites than either PB or TUDCA individually. Most changes were unique to Combo and affected the expression of genes involved in ALS-relevant pathways, such as nucleocytoplasmic transport, unfolded protein response, mitochondrial function, RNA metabolism, and innate immunity. Weighted gene coexpression network analysis showed that significant correlations between ALS gene expression modules and clinical parameters were abolished by Combo. This study is the first to explore the molecular effects of Combo in ALS patient-derived cells. It shows that Combo has a greater and distinct impact compared to the individual compounds and provides clues to drug targets and mechanisms of actions, which may underlie the benefits of this investigational drug combination.

## Introduction

ALS is a progressive and fatal neurodegenerative disorder that affects approximately every 5 per 100,000 people in the United States (1). The disease involves degeneration of both upper and lower motor neurons, causing muscle weakness and atrophy, spasticity, dysphagia, and neurocognitive symptoms. Eventual paralysis and death due to respiratory insufficiency typically occurs within 2-5 years of diagnosis (2). Approximately 30 genes have been identified as major risk factors for ALS, and 10-15% of ALS cases are associated with a causative underlying mutation in one or more of these genes, termed familial ALS (fALS). The remaining 85-90% of cases are sporadic (sALS) and may arise from a combination of genetic predispositions and environmental factors but have no defined underlying specific gene mutation (3).

Diverse pathophysiological mechanisms have been proposed to contribute to motor neuron degeneration in ALS, including gliosis/inflammation, hyperexcitability, oxidative stress, disruptions in proteostasis, endoplasmic reticulum stress, alterations in RNA processing, abnormal stress granule dynamics, impaired vesicular transport, mitochondrial dysfunction, impairment in DNA damage repair, and alterations of nuclear transport (2). The pathophysiology of ALS likely results from a combination of these mechanisms, which converge to cause motor neuron degeneration. The lack of a clear genetic cause for sALS precludes the development of representative animal models. This, together with the potential heterogeneity in etiology, has thus far hindered the development of successful therapies. The only FDA-approved drugs for ALS are Edaravone, which reduces oxidative stress, and Riluzole, which reduces excitotoxic cell death (3). The clinical efficacy of these treatments is modest, and there is an urgent need for new treatments with greater ability to slow functional decline and extend survival.

Recently, the combination of taurursodiol (also known as tauroursodeoxycholic acid, TUDCA, Turso) and sodium phenylbutyrate (PB) was investigated in a double-blind, randomized, multisite, placebo-controlled phase 2 clinical trial (CENTAUR). 137 ALS patients were randomized 2:1 to receive active drug or placebo for 24 weeks. The trial successfully hit its primary endpoint, rate of change in the ALS Functional Rating Scale (ALSFRS-R) score, with a 1.24 point/month decline in the treatment group vs. a 1.66 point/month decline in placebo (p=0.03) (4). Importantly, long-term analysis showed that the median survival was 25 months in the original treatment group compared to 18.5 months in the original placebo group, an increase of 6.5 months (5). This outcome outperforms the currently approved treatments for ALS.

TUDCA is a hydrophilic secondary bile acid produced by conjugation of taurine to ursodeoxycholic acid. Bile acids are endogenously produced in the liver from cholesterol and can also be synthesized in the brain (6). Bile acids, including TUDCA, can signal through a variety of plasma membrane and nuclear receptors including FXR, TGR5/GBPAR1, S1PR2, GR, MR, and α5b1 integrin, activating diverse signaling pathways such as Nf-kB, MAPK, PI3K, ERK, PLCg, PKA, and Akt (6–8). TUDCA can act as a chemical chaperone and alleviate ER stress due to misfolded proteins (9). TUDCA can block apoptosis by enhancing mitochondrial function, including augmenting inner mitochondrial membrane integrity, reducing reactive oxygen species (ROS) production, and increasing oxidative phosphorylation (10). Additionally, it modulates epigenetics by reducing the activity and/or expression of histone deacetylases (HDACs) and histone acetyltransferases (HATs) (6). TUDCA has showed anti-inflammatory effects in several animal and cell models of neurodegeneration (7, 11).

PB is an aromatized fatty acid that is metabolized into phenylacetate through β-oxidation, which can be then conjugated to glutamine to form phenylacetylglutamine, which acts as an ammonia sink. PB is approved by the FDA for the treatment of urea cycle disorders (12). PB is an HDAC inhibitor and modulates chromatin remodeling and transcription by increasing histone acetylation (13, 14). PB is a chemical chaperone and is protective in encephalopathies caused by protein instability (15). Evidence shows that PB can ameliorate ER stress and modulate the unfolded protein response (UPR) by preventing misfolded protein accumulation in the ER lumen and regulating protein trafficking, thereby preventing apoptosis (16). Furthermore, in the SOD1 *G93A* mouse model of fALS, PB improves survival, motor function, and histone acetylation, and reduces motor neuron loss, gliosis, ubiquitin-positive aggregates, and apoptosis (17).

TUDCA and PB are each highly multi-functional molecules, and the breadth of cellular processes targeted by these two drugs could be particularly beneficial in ALS. Additionally, their partially overlapping mechanisms of action mean that administrating them together may lead to synergistic activity. While each molecule individually has been the subject of several studies, the molecular effects of the TUDCA-PB combination have not been investigated. In this work, we investigated the processes affected by the combination of TUDCA and PB (Combo) and compare the effects of the Combo to those of each drug individually in cell lines from sALS patients and healthy controls. To this end, we performed unbiased metabolomics and transcriptomics on primary skin fibroblasts. While skin fibroblasts do not degenerate in ALS, sALS fibroblasts have been shown to manifest metabolic and transcriptomic alterations (18–22). sALS fibroblasts display altered age-related metabolic profiles and bioenergetic states compared to control fibroblasts (23, 24). Furthermore, sALS fibroblasts show pathological changes similar to those seen in disease-relevant cell types, including increased susceptibility to DNA damage, TDP-43 cytosolic mislocalization (25, 26), and oxidative phosphorylation impairment (27). Moreover, perivascular fibroblasts from sALS patients exhibit transcriptomic alterations (28). Overall, fibroblasts represent a valuable and accessible platform to study the effects of investigational therapeutics on disease-relevant pathways.

We treated fibroblasts from sALS patients and healthy controls with TUDCA, PB, and the combination of both drugs (Combo). Combo treatment caused many more changes in metabolism and gene expression than each individual drug and the effects of Combo were largely distinct from those of each drug alone. The effects of Combo encompassed changes in genes involved in mitochondrial function, UPR, intracellular trafficking/nucleo-cytoplasmic transport, innate immune function, nucleic acid metabolism, and RNA processing. While some of these pathways are shared between sALS and CTL, others are uniquely affected in each group. These analyses provide new insight into the mechanisms of action of TUDCA and PB combination in sALS patient-derived cells.

## Methods

### Cell Culture and Drug Treatment

Twelve primary fibroblast lines from healthy donors and 12 sALS patient lines were used in this study. Demographic and clinical data from de-identified subjects are shown in Table S1. Cells were cultured in Dulbecco’s modified Eagle medium (DMEM) containing 5 mM glucose, 2 mM L-glutamine, 1 mM sodium pyruvate, 10% FBS, and 1% penicillin/streptomycin. For all experiments, lines were assessed at passage 8-10. Treatment was done with 10 μM TUDCA and 100 μM PB for 5 days, replacing the medium with drug-containing media daily. Doses are within the range of observed plasma concentration of the drug in patients according to unpublished data generated by Amylyx Pharmaceuticals.

### RNA Sequencing

Cells were harvested in Trizol (Thermo Fisher), homogenized through an 18g syringe, extracted with chloroform, and centrifuged at 12,000 x g for 15 minutes at 4°C. Total RNA was isolated using the SV Total RNA Isolation System (Promega). The Genomics Facility at the Cornell Institute of Biotechnology used 500ng of RNA/sample for 3’RNA library preparation with the Lexogen QuantSeq 3’ mRNA-Seq Library Prep Kit FWD (Illumina), sequenced libraries on an Illumina NextSeq500 sequencer (single end 1×86bp), and de-multiplexed based upon six base i7 indices using Illumina bcl2fastq2 software (version 2.18). Illumina adapters were removed using Trimmomatic (version 0.36). Trimmed reads were aligned to the human genome assembly GRCh38.p13 using the STAR aligner version 2.7.0f, and transcriptome assembly was done using HTSeq-count version 0.6.1 (29).

### Metabolomics

Metabolites were rapidly extracted in 80% ice-cold methanol. Extracted samples were vortexed twice, cleared by centrifugation at 14,000 x g for 20 minutes at 4°C, and stored at −80°C. The Weill Cornell Medicine Meyer Cancer Center Proteomics & Metabolomics Core Facility performed hydrophilic interaction liquid chromatography-mass spectrometry (LC-MS) for relative quantification of polar metabolite profiles for both targeted and untargeted metabolites. Metabolites were measured on a Q Exactive Orbitrap mass spectrometer (Thermo Scientific), coupled to a Vanquish UPLC system (Thermo Scientific) via an Ion Max ion source with a HESI II probe (Thermo Scientific). A Sequant ZIC-pHILIC column (2.1 mm i.d. × 150 mm, particle size of 5 μm, Millipore Sigma) was used for separation. The MS data was processed using XCalibur 4.1 (Thermo Scientific) to obtain the metabolite signal intensity for relative quantitation. Total protein, determined by BCA Assay, was used for normalization. Targeted identification was available for 167 metabolites based on an in-house library established using known chemical standards. The remaining 629 untargeted metabolites were assigned one or more identifiers based on their mass/charge ratio, and for clarity in figures are represented with the most commonly cited metabolite identifier, with full identifier lists available in Table S1. Identification required exact mass (within 5ppm) and standard retention times.

### Data Analysis

The R package DESeq2 version 1.24.0 (30) was used for normalization and differential expression analysis, with a low counts filter of <96 and all other filtering parameters kept as defaults. A Wald test was used to determine statistical significance, with the cutoff being a False Discovery Rate <5% after Benjamini-Hochberg correction. Weighted gene co-expression network analysis was done using the normalized gene expression data from DESEq2 as input for the functions included in the WGCNA package available from CRAN (31). We optimized parameters to maintain scale-free topology, and to allow for direct comparisons between the two networks, Q-Q scaling was performed such that the 95% quantiles of both matrices matched. For all networks the module merging parameter was kept consistent at 80%. Pairwise Pearson’s correlations were used to calculate associations between modules and disease traits. Pathway analysis for all gene expression data was performed with the gprofiler2 (32) and clusterProfiler packages (33), and the cutoff for significance was a FDR corrected p-value < 0.05. Pathways shown in the figures were condensed using the simplify function from the clusterProfiler package to merge terms with more than 40% overlapping annotated genes.

Relative metabolite abundance data were normalized with a log transformation, and differential abundance and pathway analyses were done with the free online tool MetaboAnalyst 5.0 (34). Metabolite significance was determined with one-way ANOVA with post-hoc t-tests, with the cutoff being a raw p-value <0.05, and the pathway significance cutoff was a FDR corrected p-value <0.05. Multi-omics analysis was done with the mixOmics package (35). Data visualization was done in R using the ggplot2, pheatmap, corrplot, and venndiagram packages available from CRAN, and in GraphPad Prism version 9.3 (GraphPad Software, Inc). Z scores were calculated from normalized counts for each gene using the standard formula (x-μ)/σ, where x is the sample value, μ is the population mean, and σ is the population standard deviation.

## Results

### Combo has a greater and distinct effect on metabolism than either PB or TUDCA alone

Initial experiments examined the global effects of TUDCA, PB, and Combo in primary human skin fibroblasts. We utilized de-identified fibroblasts from sALS patients and age- and sex-matched healthy controls (n=12/group). Data describing patient characteristics is shown in Table 1. We performed unbiased metabolomics on cells maintained in media containing physiological glucose levels (5mM) and identified 167 targeted and 631 untargeted polar metabolites (Table S1). Partial least squares-discriminant analysis (PLS-DA) of the 798 metabolites showed that all samples (sALS and CTL) treated with Combo were largely separable from the other treatments (Fig 1A). Differential metabolite analysis identified 27 significantly different metabolites in Combo treated samples vs. vehicle (PBS) treated samples, with only 10 and 8 significant metabolites with TUDCA and PB, respectively (Fig 1B, Table S2). The majority (25/27) of significant metabolites in Combo were unique to this treatment (Fig 1C). Therefore, Combo has distinct effects on metabolism that are not simply additive effects of PB and TUDCA.

**Figure 1.**
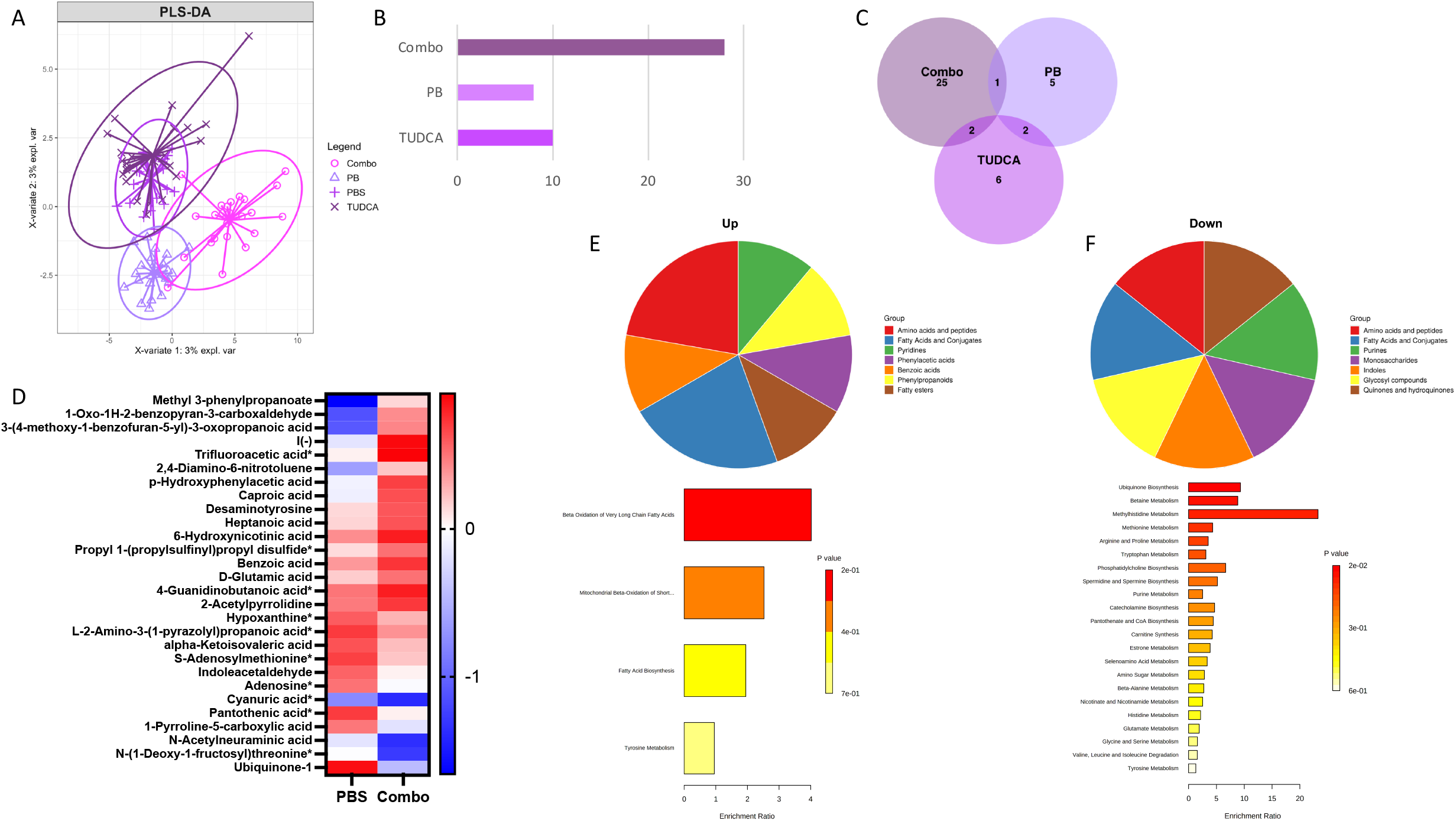
Combo changes more metabolites than either PB or TUDCA alone. A) Partial least squares-discriminant analysis of normalized abundance data for all metabolites. B) Bar graph showing number of significantly different metabolites (p-value < 0.05) for each treatment compared to vehicle. C) Venn diagram of metabolites significantly changed by each treatment (p-value < 0.05). D) Heatmap of Z-scores of all metabolites significantly changed by Combo. * Indicates targeted metabolites. E-F) SMPDB category and pathway analysis of Combo upregulated (E) and downregulated (F) metabolites.

**Table 1:** Demographic and clinical characteristics of de-identified fibroblast lines used in this study. * % Calculated as ((48 - ALSFRS at Skin BX) / Disease Duration [months] at Skin BX)

|  | ID | Age at biopsy | Sex | Disease duration at time of skin biopsy (months) | ALSFRS-R total at biopsy | Rate of ALSFRS-R decline* | FVC % |
| --- | --- | --- | --- | --- | --- | --- | --- |
| **Control** | 1 | 55 | F |  |  |  |  |
|  | 2 | 59 | M |  |  |  |  |
|  | 3 | 69 | M |  |  |  |  |
|  | 4 | 54 | M |  |  |  |  |
|  | 5 | 65 | F |  |  |  |  |
|  | 6 | 61 | F |  |  |  |  |
|  | 7 | 79 | M |  |  |  |  |
|  | 8 | 53 | F |  |  |  |  |
|  | 9 | 58 | M |  |  |  |  |
|  | 10 | 65 | M |  |  |  |  |
|  | 11 | 62 | F |  |  |  |  |
|  | 12 | 51 | M |  |  |  |  |
| **sALS** | 1 | 64 | M | 14 | 35 | 0.93 | 79 |
|  | 2 | 77 | M | 8 | 30 | 2.25 | 83 |
|  | 3 | 67 | F | 14 | 35 | 0.93 | 78 |
|  | 4 | 64 | M | 24 | 38 | 0.42 | 55 |
|  | 5 | 41 | F | 9 | 30 | 2 | 90 |
|  | 6 | 54 | F | 10 | 22 | 2.6 | 39 |
|  | 7 | 64 | F | 15 | 28 | 1.33 | 22 |
|  | 8 | 60 | F | 10 | 41 | 0.7 | 115 |
|  | 9 | 64 | M | 15 | 27 | 1.4 | 45 |
|  | 10 | 64 | F | 8 | 36 | 1.5 | 86 |
|  | 11 | 68 | F | 12 | 24 | 2 | 98 |
|  | 12 | 65 | M | 11 | 28 | 1.82 | 47 |

Both targeted and untargeted metabolites changed by Combo are shown in Fig 1D. In Combo-treated samples, upregulated metabolites included several fatty acids, including caproic acid and heptanoic acid, and methyl-3-propanoate, although we note that the list of potential identifiers assigned to this metabolite included a PB derivative, suggesting that this increase is partly due to drug metabolism. The most strongly downregulated metabolite was ubiquinone-1. Pantothenic acid (Vitamin B5), adenosine, s-adenosylmethionine, and hypoxanthine (a product of purine breakdown) were also downregulated. The major metabolic pathways upregulated by Combo treatment were related to fatty acid oxidation, and when upregulated metabolites were grouped according to their Small Molecule Pathway Database (SMPDB) class, the majority were amino acids and fatty acids (Fig 1E). Downregulated metabolites were evenly split across several SMPDB classes, and approximately half the significantly enriched pathways (12/22) were related to amino acid metabolism (Fig 1F).

### Combo has a greater and distinct effect on gene expression in fibroblasts than either PB or TUDCA alone and regulates the expression of ALS-relevant genes

Next, we performed unbiased 3’ RNA sequencing on total RNA. We found that the major contributor to overall variability in gene expression among fibroblast lines was inter-line variability. Principal component analysis (PCA) of the top 500 most variable genes in all lines showed that the factor driving clustering was the individual line identifier (Fig 2A). Samples from the same cell line clustered tightly together, regardless of treatment, and no clear clustering due to treatment was apparent. However, PLS-DA of the top 500 most variable genes showed that while PB and TUDCA treated samples fully overlapped with PBS treated samples, Combo treated samples could be separated (Fig 2B). Therefore, Combo induced greater changes in overall gene expression than PB or TUDCA alone, similar to what was found in metabolomic analysis. Differential expression analysis showed that fewer differentially expressed genes (DEGs), defined as having an adjusted p-value < 0.05, were found in the TUDCA vs. PBS comparison (16 DEGs, Fig 2C) and PB vs. PBS (186 DEGs, Fig 2D) when compared to Combo vs. PBS (1838 DEGs, Fig 2E, Table S3). Similar to metabolites, the vast majority (1796/1838) of DEGs in the Combo vs. PBS were unique to Combo treatment (Fig 2F).

**Figure 2.**
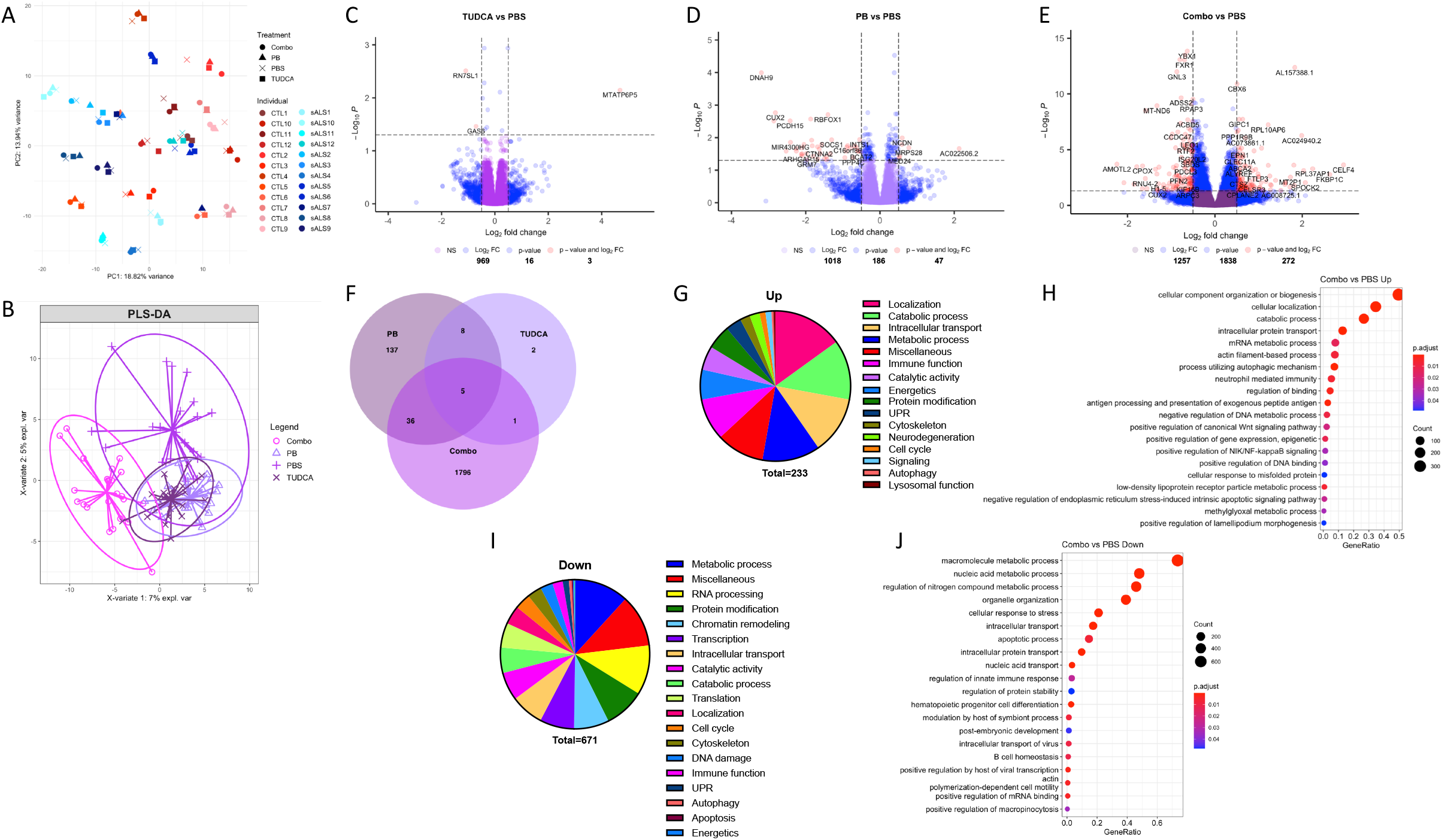
Combo changes more genes than either PB or TUDCA alone. A) Principal component analysis of the top 500 genes with highest variability. B) Principal least squares-discriminant analysis of the top 500 genes with highest variability. C-E) Volcano plots of DEGs of TUDCA (C), PB (D), and Combo (E) relative to vehicle. F) Venn diagram of DEGs in each treatment. G) Category analysis of all significant GO terms enriched in Combo upregulated DEGs. H) Top 20 most significant GO:BP terms enriched in Combo upregulated DEGs, simplified with the clusterProfiler package. I) Category analysis of all significant GO terms enriched in Combo downregulated DEGs. J) Top 20 most significant GO:BP terms enriched in Combo downregulated DEGs.

We performed Gene Ontology (GO) analysis using annotated pathways from the GO Molecular Function (GO:MF), GO Biological Process (GO:BP), and Kyoto Encyclopedia of Genes and Genomes (KEGG) databases to look for processes and categories enriched among the DEGs. Numerous pathways were upregulated by Combo (233 total, Table S4), with one of the most significantly enriched pathways was “cellular protein localization” (adjusted p-value= 2.37e-13) (Fig 2H). The largest proportion of enriched pathways was related to macromolecule localization and transport, including “endocytosis” (p_adj_=0.009), “exocytosis” (p_adj_=0.001), “endosomal transport” (p_adj_ = 0.028), “golgi vesicle transport” (p_adj_=0.049), and “nuclear transport” (p_adj_=0.028) (Fig 1G). Nuclear transport alterations are known to occur in ALS (36), and we found several genes belonging to the importin (*TNPO3, IPO5, IPO7*) and nuclear pore complex (*TPR, NUP188, NUP153, NUP54, POM121*) changed by Combo treatment (Fig S1A). Another major group of pathways associated with ALS found upregulated by Combo treatment included those related to_bioenergetics, such as “aerobic respiration” (p_adj_=0.019), “ATP synthesis coupled electron transport” (p_adj_=0.04), “inner mitochondrial membrane organization” (p_adj_=0.012), and “oxidative phosphorylation” (p_adj_=0.008). One or more genes encoding subunits of each mitochondrial oxidative phosphorylation complex (except for Complex II) were upregulated in Combo-treated cells (Fig S1B). Furthermore, another group of pathways relevant to ALS pathogenesis was related to the UPR, including “cellular response to misfolded protein” (p_adj_=0.049), “ERAD pathway” (p_adj_=0.008), “response to endoplasmic reticulum stress” (p_adj_=0.004), and “protein quality control for misfolded or incompletely synthesized proteins” (p_adj_=0.023) (Fig 1G). Upregulated UPR genes included *WFS1* and *CREB3*, both of which promote cell survival under ER stress (37, 38). Of note, Combo is also being studied for the treatment of Wolfram Syndrome, which is caused by mutations in *WFS1* (39). *VCP*, a segregase that recruits ubiquitinated misfolded proteins to the proteasome for degradation and causes fALS when mutated (40), was also upregulated by Combo (Fig S1C). Interestingly, we also found several pathways associated with innate immune activation, and in particular genes encoding for subunits of the 20S (*PSME2, PSMD3, PSMD4*) and 26S (*PSMB5, PSMB6*) immunoproteasomes upregulated by Combo (Fig S1D). The immunoproteasome degrades antigens presented by MHC-I, and may also be important for the clearance of misfolded proteins (41). Furthermore, we found STING1 upregulated and cGAS downregulated by Combo (Fig S1D), and the cGAS/STING1 axis of innate immune activation has recently been shown to be the consequence of mitochondrial dysfunction in fALS (42). Overall, these findings confirm the upregulation by Combo of pathways associated with the mechanism of action (MOA) of TUDCA and PB (9, 10, 16) such as alleviating mitochondrial dysfunction and ER stress, but also highlight other possible disease-relevant pathways affected by Combo.

One of the largest categories of pathways downregulated by Combo was related to RNA metabolism and processing, a major interest in ALS (43). Some of the most significantly enriched downregulated pathways included “RNA metabolic process” (p_adj_=9.11e-30), “nucleic acid metabolic process” (p_adj_=3.26e-33), and “RNA processing” (p_adj_=3.57e-29) (Fig 1I). Additional terms related to RNA processing included “RNA splicing” (p_adj_=3.87e-19), “RNA stabilization” (p_adj_=0.003), “RNA transport” (p_adj_=2.51e-7), and “spliceosome” (p_adj_=0.001) (Fig 1H, Table S4). Enriched terms encompassed processing of mRNA, rRNA, ncRNA, and miRNA, and include polyadenylation, capping, and methylation activities. Other downregulated pathways were related to protein modifications, including acetylation, methylation, and polyubiquitination. Many of the genes in these pathways are also associated with chromatin remodeling, such as “histone acetylation” (p_adj_=0.003), “histone methylation” (p_adj_=0.0001), “histone ubiquitination” (p_adj_=0.01), and “nucleosome assembly” (p_adj_=0.003). Specific histone modification genes identified in downregulated pathways encode for enzymes that perform H2B ubiquitination and H3 methylation at K36 and K4, marks associated with actively transcribed chromatin (44). Downregulated transcription-associated pathways included “transcription by RNA polymerase II” (p_adj_=4.35e-6), the major polymerase involved in transcribing mRNAs and miRNAs. Terms were related to phosphorylation of RNA Pol II, transcriptional initiation, and both positive and negative regulators of the polymerase. Genes identified in these pathways included members of the mediator transcriptional co-activator complex (*MED4, MED13, MED19*, and *MED29*) (45), components of the basal transcription factor TFIID (*TAF1, TAF4B*) (46), *GLYR1*, which enables RNA Pol II transcription by destabilizing nucleosomes (47), *YBX1*, a transcription factor that regulates oncogenic and immune genes (48), and *LEO1* and *TCEA1*, both important for RNA Pol II transcriptional elongation (49, 50). Another category of downregulated pathways relevant to ALS was DNA damage repair, including “cellular response to DNA damage stimulus” (p_adj_=1.64e-11), “double-strand break repair” (p_adj_=0.0006), “nucleotide-excision repair” (p_adj_=0.002), and “recombinatorial repair” (p_adj_=0.016). Taken together, GO analysis reveal a global downregulation of genes involved in a broad range of cellular processes, including transcription and RNA metabolism and processing.

To better understand the transcriptional regulation underlying Combo-driven gene expression changes, we performed transcription factor binding site enrichment analysis on DEGs up- and downregulated by Combo. The largest groups of transcription factor binding sites enriched in both up- and downregulated genes belonged to Elk1, E2F1, and Elf1, while Sp1 and Egr1 were selectively enriched in upregulated genes, and YY1 and CREB1 in downregulated genes (Table S5). Elk1, E2F1, and Egr1 have important roles in regulating the cell cycle and apoptosis (51–53). Sp1 controls a broad range of cellular functions, including cell survival, immune responses, DNA damage responses, and chromatin remodeling (54). Elf1 is an important regulator of the immune response (55), and YY1 modulates DNA damage repair (56). The functions of these transcription factors align with the GO pathways identified in Combo-regulated genes (Fig 2), suggesting that these transcription factors are effectors of Combo treatment.

### Multi-omics analysis identifies a set of 5 metabolites and 20 genes sufficient to discriminate Combo treatment from vehicle treatment

Next, we combined metabolomic and transcriptomic datasets for multi-omics analysis to identify related features of both datasets able to distinguish Combo from the other treatments. We used the DIABLO (Data Integration Analysis for Biomarker discovery using Latent variable approaches for Omics studies) algorithm found by PLS-DA that Combo-treated cells (sALS and CTL combined) could be partially separated from all other treatments on Variate 1, which utilized the most discriminatory 5 metabolites and 20 genes (Fig 3A). Receiver-operating characteristic (ROC) curves of a classification model using these metabolites and genes showed that discriminating Combo-treated samples from all other samples performed better than any other classification, confirming that Combo treatment had a more distinct effect on gene expression and metabolism than either drug individually (Fig 3B). To further examine the genes and metabolites driving the classification of Combo from other groups, we performed the analysis using only Combo and PBS samples. PLS-DA identified the most discriminatory 5 metabolites and 20 genes and showed that Combo could be fully separated from PBS on Variate 1 (Fig 3C). Hierarchical clustering confirmed full separation of Combo from PBS based on the Variate 1 metabolites and genes (Fig 3D). Several of the 20 genes that most strongly discriminated Combo from PBS samples were RNA binding genes involved in RNA polymerase II transcription, including *MED4, FXR1, YBX1*, and *GLYR1* (45, 47, 48, 57). Expression of these genes was decreased after Combo. The metabolites that most strongly discriminated Combo from PBS included methyl-3-propanoate/3-phenylbutyric acid and ubiquinone-1, and also I(-), or hydroiodic acid. Taken together, the combined multi-omics (Fig 3) and individual dataset analyses (Fig 1-2) confirmed that Combo has unique effects on gene expression and metabolism, strongly driven by a subset of genes and metabolites identified with both methods.

**Figure 3.**
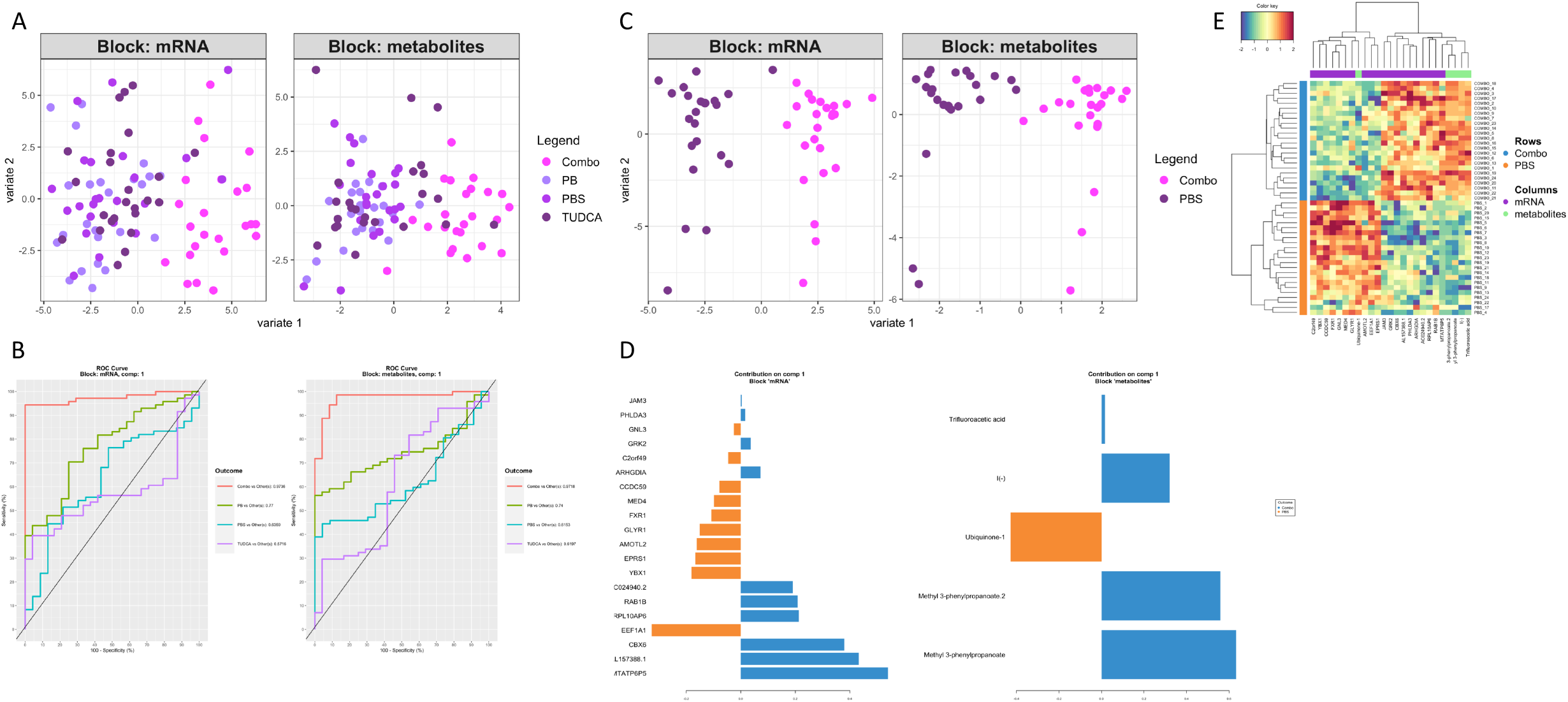
A set of 20 genes and 5 metabolites can discriminate Combo-treated samples from vehicle. A) Principal least squares-discriminant analysis of multi-omics comparison normalized RNA and metabolites for all treatments, based on the top 20 most discriminatory genes and top 5 most discriminatory metabolites. B) ROC curves for the discrimination of each treatment from all other treatments based on the same 20 genes and 5 metabolites as in (A). C) Principal least squares-discriminant analysis of multi-omics comparison normalized RNA and metabolites for Combo vs.PBS based on the top 20 most discriminatory genes or top 5 most discriminatory metabolites. D) Bar graphs of the top 20 genes and top 5 metabolites for Combo vs. PBS. E) Hierarchical clustering of samples based on the values of the 20 genes and 5 metabolites as in (C-D).

### Combo has a greater and distinct effect on gene expression of sALS compared to CTL fibroblasts

We then analyzed CTL and sALS cell lines separately to determine whether the effects of Combo treatment are different in these two groups. All metabolites significantly changed by Combo in sALS were also changed in CTL (Fig S2). On the other hand, there were double the number of statistically significant DEGs in sALS relative to CTL (522 vs. 223) (Fig 4A-B, Table S6-7). Half of the upregulated DEGs in the CTL Combo vs. PBS comparison overlapped with those identified in sALS (33/66) and approximately ~66% of the downregulated DEGs also overlapped (96/157), the majority of DEGs in the sALS Combo vs. PBS comparison were unique to this group (Fig 4C). The same pattern was evident for significantly enriched pathways (Fig 4D). While we found only 4 pathways significantly enriched in upregulated DEGs in CTL Combo vs. PBS, all related to RNA splicing (Fig 4E, Table S8), there were 67 enriched pathways in the sALS Combo vs. PBS upregulated DEGs (Table S9). Major categories of pathways upregulated by Combo in sALS included intracellular transport, cytoskeleton organization, and autophagy (Fig 4F-G). The strong enrichment of intracellular localization and transport pathways matched the enrichment findings from the sALS and CTL combined analysis (Fig 2), suggesting that the enrichment of these pathways is driven by sALS lines. Within the intracellular transport category, many of the DEGs identified in the combined analysis (Table S5) overlapped with those found specifically in sALS (Table S10). Within the cytoskeleton organization category, we found regulators of Rho GTPases and vesicle trafficking, and actin binding proteins to be upregulated (Table S10). In the autophagy category, we found regulators of autophagosome formation, including *GRAMD1A* (an ER cholesterol transporter) (58), *RAB1B* (an ER GTPase) (59), *PACS2* (a mitochondrial-ER contact protein) (60), and mTORC1 regulator *LAMTOR4* (61). All of these, except *RAB1B*, were uniquely upregulated in sALS. These findings are interesting because both TUDCA and PB were shown to modulate cellular responses to ER stress (9, 10, 16).

**Figure 4.**
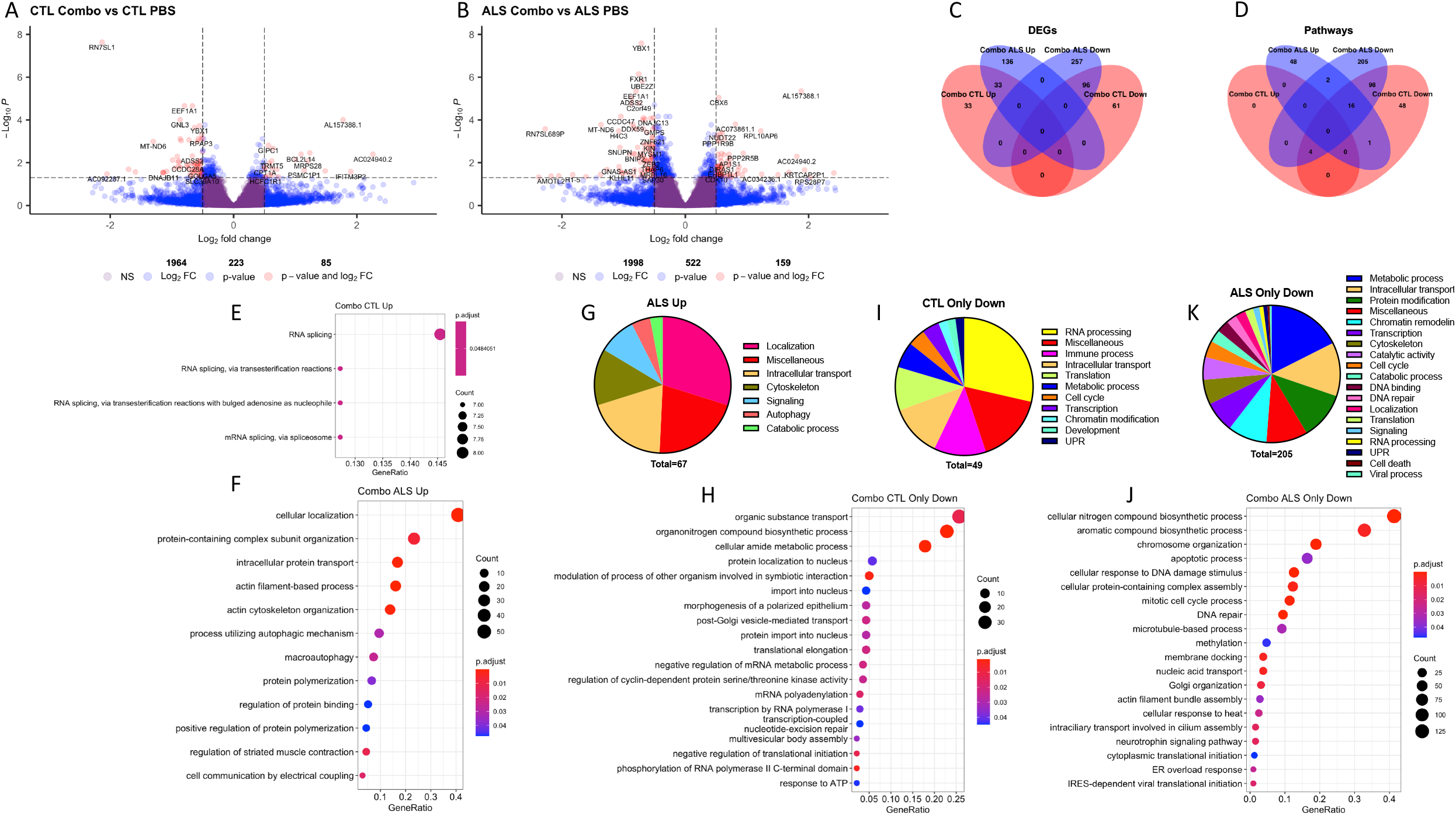
Combo has different transcriptional effects in sALS and CTL cells. A-B) Volcano plots of DEGs in the Combo vs. PBS comparison in CTL (A) and sALS (B) lines. C-D) Venn diagram comparing DEGs (C) and GO terms (D) changed by Combo in sALS and CTL lines. E) Top 4 most significant GO:BP terms enriched in Combo upregulated DEGs in CTL. F) Top 20 most significant GO:BP terms enriched in Combo upregulated DEGs in ALS. G) Category analysis of all significant GO terms enriched in Combo upregulated DEGs in ALS. H) Top 20 most significant GO:BP terms enriched in Combo downregulated DEGs in CTL. I) Category analysis of all significant GO terms enriched in Combo downregulated DEGs in CTL. J) Top 20 most significant GO:BP terms enriched in Combo downregulated DEGs in ALS. K) Category analysis of all significant GO terms enriched in Combo downregulated DEGs in ALS.

We next identified pathways specifically downregulated by Combo in CTL and sALS (Table S8-9). The largest category of terms uniquely downregulated by the treatment in CTL was RNA processing. Most of these terms were associated with RNA binding, splicing, and polyadenylation (Fig 4H-I). Uniquely in sALS, the largest category of downregulated terms was metabolic processes, mainly related to nucleic acid metabolism (Fig 4J-K). Furthermore, more genes and terms related to RNA polymerase II transcription were downregulated by Combo treatment in sALS compared to CTL. These include *MED19* and *TAF2*, both involved in RNA polymerase II transcription elongation (45, 47), and *SETX*, a helicase that modulates RNA polymerase II binding to chromatin, and interestingly a genetic cause of juvenile ALS (62).

Taken together, these GO results reveal that many of the transcriptional effects of Combo are unique to sALS. In particular, Combo modulates the expression of intracellular transport and RNA polymerase II transcription genes only in sALS, both of which may be relevant to disease pathogenesis.

### WGCNA identifies module-trait associations in vehicle-treated cells which are modified by Combo treatment

We extended our analysis of the effects of Combo on gene expression to include weighted gene co-expression network analysis (WGCNA). WGCNA can complement differential expression analysis methods. Rather than using statistical significance based on differing levels of expression between groups as a cutoff, WGCNA considers highly similar groups of gene based on expression patterns across samples as sets of interconnected modules (31, 63), thereby creating a network of gene expression patterns which can then be correlated to disease traits. This approach increases statistical power for identifying associations between traits and gene expression profiles. WGCNA has recently been applied to discover transcriptomic alterations in several neurodegenerative diseases, including ALS (64).

We used normalized gene expression data from 16,492 genes that passed quality control filters as input to construct two co-expression networks, one from all 24 Combo-treated samples, and the other from all 24 vehicle (PBS)-treated samples (Fig 5A-B). After hierarchical clustering to identify groups of highly co-expressed genes (modules), we found 49 modules in the Combo network and 40 modules in the vehicle network (Fig 5C-D). We then correlated module gene expression with disease traits, including disease state (sALS vs. CTL), disease duration (time between symptom onset and skin biopsy), ALS Functional Rating Score-Revised (ALSFRS-R) score, rate of decline in ALSFRS-R, and forced vital capacity (FVC)%, as well as individual cell line number, age, and sex. We included age and sex as relevant traits for correlation because the incidence of ALS varies with both, making them potential biologically relevant variables. In vehicle-treated samples, sex was not a major driver of variance in overall gene expression, as it did not correlate with any of the first three principal components (PCs), while age significantly correlated with PC1 (Fig S5). In Combo-treated samples, sex and age did not correlate with any of the top three PCs (Fig S5). Therefore, variance in gene expression associated with age that exists in vehicle-treated cells was attenuated by Combo treatment.

**Figure 5.**
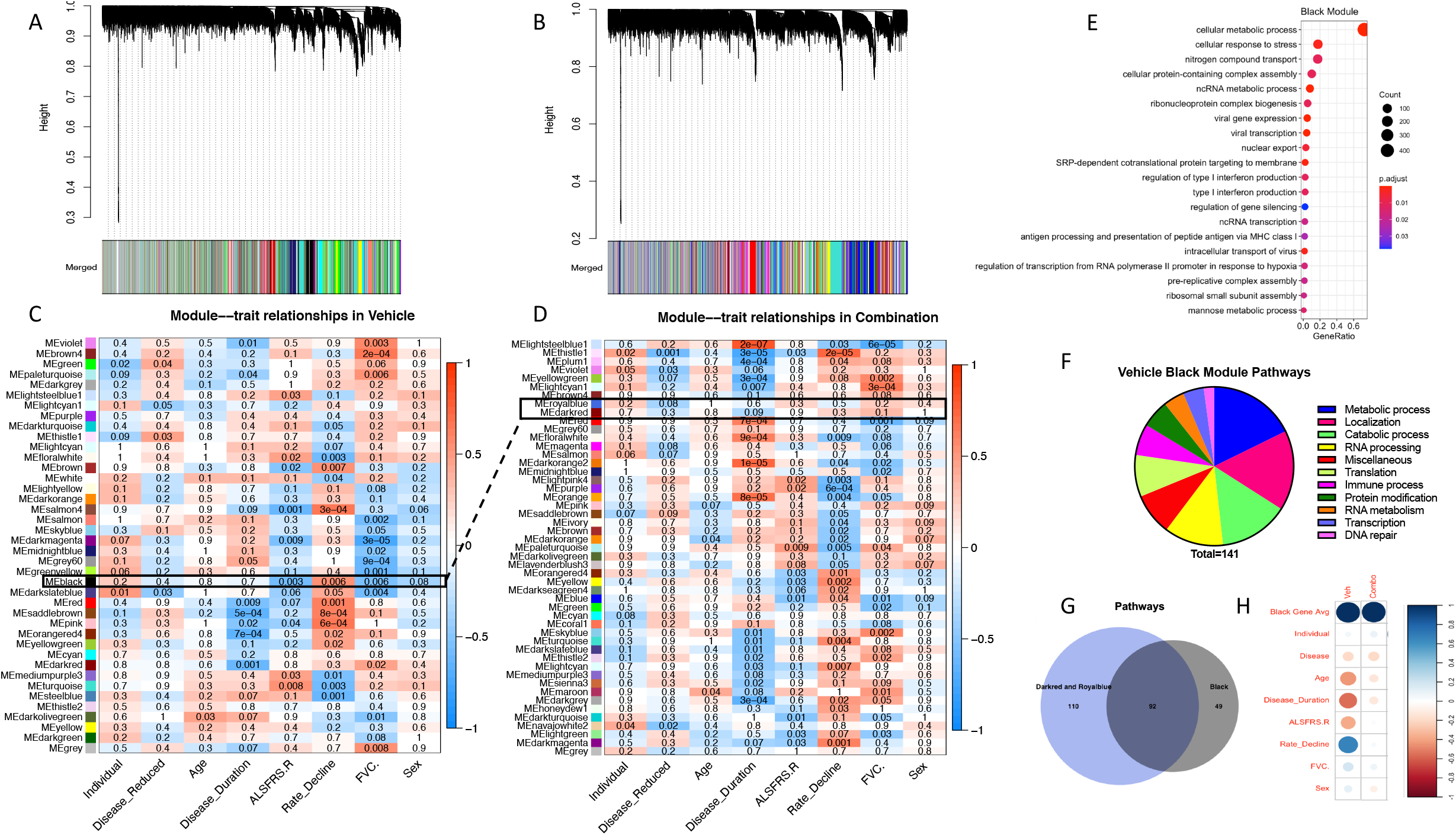
WGCNA identifies modules of genes associated with ALS clinical traits that are altered by Combo. A-B) Dendograms of hierarchical clustering of gene co-expression modules for vehicle-treated samples (A) and Combo-treated samples (B). C-D) Heatmaps showing correlations between module eigengene expression values and clinical traits for vehicle-treated samples (C) and Combo-treated samples (D). The numbers in each box are p-values, and box colors correspond to the correlation coefficient. E) Top 20 most significant GO:BP terms enriched in the vehicle Black module. F) Category analysis of all significant GO terms enriched in the vehicle Black module. G) Venn diagram of overlap between significantly enriched GO pathways found in the vehicle Black module and the Combo modules (Darkred and Royalblue) paired with Black. H) Correlations between the expression of the 758 Black module genes in vehicle- and Combo-treated samples and disease traits. Size correlates inversely to p-value, and color corresponds to Pearson’s correlation coefficient.

We identified 22/40 modules that significantly (adjusted p-value < 0.01) associated with one or more traits in the vehicle network, and 23/49 modules in the Combo network (Fig 5C-D). Of the 22 trait-associated modules from the vehicle network, 15 had at least one significant GO enrichment (Table 2). Of the 23 from the Combo network, 18 had at least one significant enrichment (Table 3). To identify functional differences in disease trait-associated modules in response to Combo, we compared the GO results from modules in each network. A larger portion of the vehicle trait-associated modules, particularly Grey60, Salmon, and Violet, had enrichment for terms related to developmental processes and the cell cycle, including specific developmental signaling pathways such as Wnt (Fig S6, Table S10). These pathways were not detected in Combo-treated modules. Furthermore, several terms related to cell death caused by oxidative stress were identified in the Paleturquoise and Violet modules and were not found in any Combo-treated modules (Fig S6, Table S10). After Combo treatment, major categories of enriched pathways that emerged included immune activation, including the terms “innate immune response”, “cytokine-mediated signaling pathway”, and “type I interferon signaling pathway” in the Turquoise and Darkturquoise modules, and translation, both cytoplasmic and mitochondrial, in the Plum1 and Blue modules (Fig S6, Table S10). Within each disease trait, the majority of GO terms enriched in all modules associated with that trait were not shared between vehicle and Combo (Fig S7). Therefore, the functions of sets of genes associated with disease traits are modified by Combo treatment.

**Table 2:** Modules in the vehicle network with a significant trait association and GO annotation

| Disease | Disease Duration | ALSFRS-R | Rate of Decline | FVC |
| --- | --- | --- | --- | --- |
|  | Darkred<br>Violet<br>Red<br>Saddlebrown | Black<br>Salmon4<br>Turquoise | Black<br>Salmon4<br>Turquoise<br>Red<br>Saddlebrown<br>Floralwhite<br>Brown<br>Pink | Black<br>Violet<br>Salmon<br>Grey60<br>Darkolivegreen<br>Grey<br>Paleturquoise |

**Table 3:**
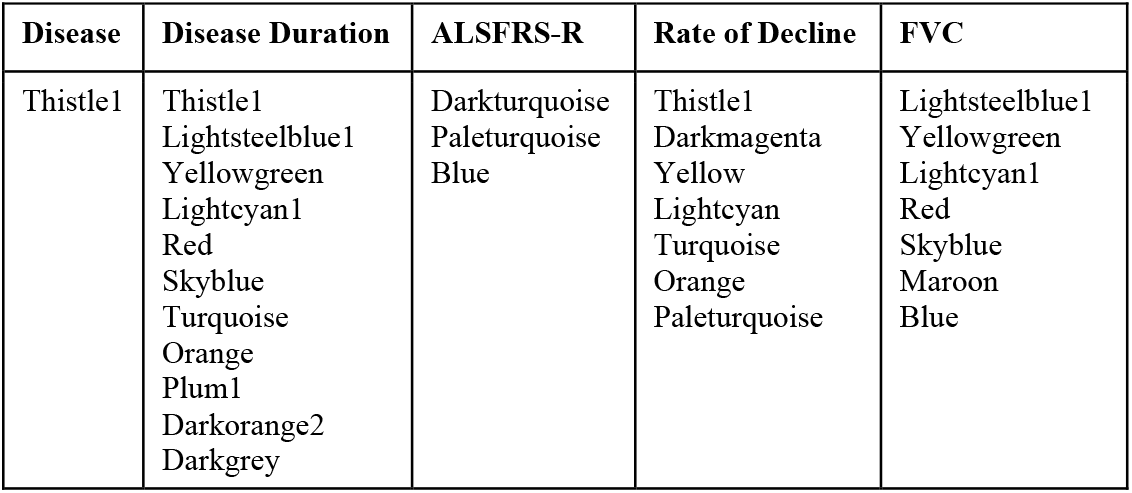
Modules in the vehicle network with a significant trait association and GO annotation

| Disease | Disease Duration | ALSFRS-R | Rate of Decline | FVC |
| --- | --- | --- | --- | --- |
| Thistle1 | Thistle1<br>Lightsteelblue1<br>Yellowgreen<br>Lightcyan1<br>Red<br>Skyblue<br>Turquoise<br>Orange<br>Plum1<br>Darkorange2<br>Darkgrey | Darkturquoise<br>Paleturquoise<br>Blue | Thistle1<br>Darkmagenta<br>Yellow<br>Lightcyan<br>Turquoise<br>Orange<br>Paleturquoise | Lightsteelblue1<br>Yellowgreen<br>Lightcyan1<br>Red<br>Skyblue<br>Maroon<br>Blue |

To further investigate changes in module-trait associations caused by Combo treatment, we compared modules from each network based on their components with Fisher’s exact test, considering two or more modules as a matched pair if adjusted p-value < 0.05 (Fig S8). 31/40 vehicle modules had a match in the Combo network. Notably, the vehicle Black module, associated with ALSFRS-R, rate of decline, and FVC%, was paired with the Combo Darkred and Royalblue modules, but neither of these had significant associations with any of these disease traits. This pair was the most interesting, because it was the only pair in which associations were lost after Combo treatment. GO analysis of the 758 genes that make up the Black module revealed a strong enrichment of RNA processing and metabolism pathways across multiple classes of RNA (ncRNA, tRNA, mt-tRNA, rRNA, and mRNA, Fig 5E). Another major class of pathways enriched in genes from the vehicle Black module was intracellular localization/transport (Fig 5F). Furthermore, individual Black module genes that had the strongest correlations with disease traits included *SELENOO*, a mitochondrial redox-sensitive selenoprotein (65), *LONP1*, a mitochondrial protease that degrades misfolded or oxidatively damaged proteins (66), *UGGT1*, which recognizes unfolded glycoproteins in the ER (67), *ORAI1*, an essential component of ER store-operated calcium entry (68), and *TRAM2* and *TMEM147*, members of the ER translocon complex (69) (Table S3). Finally, the most commonly identified transcription factors with binding sites enriched in genes from the Black module were Elk1, E2F1, Elf1, Sp1, and Erg1 (Table S11), the same transcription factors identified in the differential expression analysis. These results support and extend those obtained through differential expression analysis (Figs 2 and 4), and point to RNA processing/metabolism, intracellular transport, and ER and mitochondrial homeostasis as key ALS-related pathways modified by Combo.

Finally, as the Combo Darkred and Royalblue modules paired with the vehicle Black module (with 40/164 and 129/642 genes in common with Black, respectively, Fig S8), but did not associate with disease traits, we tested whether the loss of the association was due to functional differences in the set of non-overlapping genes from each module. We think that this is unlikely, as 92 of the 141 enriched pathways in the Black module were also found in the Darkred and Royalblue modules (Fig 5G), suggesting that there were no major functional differences. Furthermore, as the overlap between the genes in the Black module and those in Darkred and Royalblue was not complete (589/758 Black module genes not found in either Darkred or Royalblue, Fig S8), we took all 758 Black module genes and calculated correlations between their average expression in Combo-treated samples and the disease traits. This approach confirmed that the strong correlation of this set of genes with disease traits in vehicle samples was significantly reduced by Combo treatment (Fig 5H). Taken together, these results identify a set of highly coexpressed genes, correlated strongly with several measures of ALS disease severity, that are modified by Combo such that after treatment their expression no longer correlates with disease traits.

## Discussion

ALS is a rapidly progressive, fatal disease, but despite a clear need for better treatment strategies, clinical trial outcomes in ALS have been largely unsuccessful. Factors potentially driving the failure of investigational treatments include delayed diagnosis due to heterogeneity of disease presentation, a lack of disease biomarkers, and the disconnect between disease etiology in animal models of fALS and human sALS cases. The search for new treatments continues despite these hurdles, and evidence, preclinical and clinical, point to TUDCA and PB as potential therapies for ALS. TUDCA (70) and PB (71) were tested individually in Phase 2 clinical trials showing modest improvements. In the Phase 2 CENTAUR trial, the Turso-PB combination (AMX0035) significantly slowed ALSFRS-R decline (4) and extended survival in the open-label extension trial (5). Turso-PB is currently under clinical investigation in the multi-center Phase 3 PHOENIX trial, a 600-participant, randomized, 48-week clinical trial, a compassionate-use setting for CENTAUR trial participants to continue drug therapy, and a pharmacokinetic-pharmacodynamic (PK/PD) trial. While the clinical investigation of this drug combination is advancing, the molecular effects of the Combo in human cells have not yet been characterized.

In this study, we elucidate the transcriptomic and metabolic effects of the TUDCA-PB combination using unbiased approaches in primary skin fibroblasts from sALS patients and healthy controls. We compared the effects of Combo to each individual drug. Remarkably, Combo changed many more metabolites and genes than either TUDCA or PB alone, and most changes were unique to Combo treatment. Among metabolites, S-adenosylmethionine (SAM) downregulation is of particular interest. SAM can be metabolized through the transsulfuration pathway to produce cysteine, which can then be used for glutathione synthesis (72). This pathway has been shown to be altered in primary sALS patient cells (19). However, in this case other metabolites of the transsulfuration pathway, including S-adenosylhomocysteine, L-cystathione, glutathione, and oxidized glutathione, were unchanged, suggesting that the reduction in SAM did not affect this pathway significantly. On the other hand, SAM can be used as a methyl donor for histone and DNA methyltransferases, and therefore alterations in SAM levels may impact epigenetics of the cell (73). In addition to potential effects on methylation, PB is an HDAC inhibitor with known epigenetic effects (13, 14), and TUDCA can also modulate chromatin modifying enzymes (6). Numerous studies have linked changes in DNA methylation and histone post-translational modifications to ALS (74), including in a large collection of induced pluripotent stem cell-derived motor neurons (75). Furthermore, altered DNA methylation patterns were recently described in a large-scale study of blood samples from ALS patients compared to controls (76). Interestingly, we find that Combo treatment downregulated the expression of many genes involved in chromatin remodeling and others related to the transcriptional activity of RNA Pol II. Future studies investigating the epigenetic effects of Turso-PB will expand upon these findings.

Overall, we find that many ALS-relevant genes and GO pathways are changed by Combo. For example, several nucleocytoplasmic transport (NCT) pathways were altered by Combo. NCT is a process with well-validated alterations in ALS (77, 78), as several models of ALS have demonstrated abnormal nuclear membrane shape, nuclear pore complex (NPC) protein loss or mislocalization, and dysregulated NCT dynamics (79). NCT pathways were both up- and downregulated by Combo treatment, although the majority of significantly changed nucleoporin (NUP) genes were downregulated. Phenylalanine-glycine-rich NUPs are known to be mislocalized and aggregate with TDP-43 in ALS (80). Several of these NUPs were downregulated by Combo, including *Nup50, Nup54, Nup153, Nup88*, and *Nup98*. Interestingly, the transcriptional changes produced by Combo in NCT pathways were more pronounced in sALS, although the functional consequences of these changes in ALS remain to be investigated. Investigating Turso-PB in cellular and animal models of ALS with clear NCT defects may provide insight into the relevance of this pathway to drug activity and disease pathogenesis. Consistent with the known effects of TUDCA and PB against ER stress, we observe upregulation by Combo of genes promoting survival in ER stress conditions, such as *WFS1* and *CREB3*, and downregulation of *ATF6* and *EIF2AK3* (PERK), key mediators of UPR signaling (81). Furthermore, Combo upregulated expression of *VCP*, a protein whose loss of function causes fALS and other degenerative disorders (40). Combo increased the expression of several subunits of mitochondrial respiratory chain complexes, which could have beneficial effects on bioenergetic capacity, known to be impaired in both genetic and sporadic forms of ALS (82). We identified several genes connected to RNA pol II transcription strongly downregulated by Combo, including *YBX1* and *FXR1*. RNA pol II transcription-associated DNA damage is increased in ALS (83), and these two specific genes are dysregulated and affect RNA homeostasis and transcriptional activity in sALS (84, 85). Lastly, Combo modulates innate immune pathways involving cGAS/STING signaling, which has been reported to be activated in ALS (42). Subunits of the immunoproteasome were also upregulated by Combo, suggesting that Combo promotes activation of protein degradation systems, which could have a protective role in ALS (86).

In addition to differential expression analysis, we used WGCNA to correlate changes in gene expression patterns driven by Combo treatment with ALS clinical parameters, such as ALSFRS-R score and FVC%. In sALS fibroblasts, we found associations between innate immune pathways and ALSFRS-R and FVC% in both the vehicle and Combo networks. The Black module was strongly correlated with disease duration, ALSFRS-R, and FVC% in the vehicle network, and significantly enriched for several immune-related GO terms including “type I interferon production”, “interleukin-1 mediated signaling pathway”, and “viral process”. Interestingly, WGCNA using data from post-mortem ALS spinal cord samples showed a strong correlation between expression of mitochondrial oxidative phosphorylation and immune activation genes and disease (64). Importantly, in our fibroblast Combo network, the Darkred and Royalblue modules, which significantly matched as a pair with the Black module in the vehicle network, lost all associations with disease traits. The loss of these associations suggests that Combo affects the expression of inflammation-related genes, which are strongly correlated with measures of disease severity. The similarities between the findings in fibroblast and spinal cord suggest that sALS fibroblasts share common transcriptomic alterations with affected tissues from ALS patients and could provide clues to understanding the therapeutic mechanisms of action of Combo in ALS.

Many questions remain to be answered by future work. This study was done in primary fibroblasts from sALS patients, as these cells are accessible and easily manipulated to study drug effects. The extent to which the findings from fibroblasts are recapitulated in motor neurons, the primary cell type affected in ALS will need to be investigated. Furthermore, we studied drug effects at a single time point. As treatment in patients will extend into months or years, the longitudinal effects of Combo should be a focus of future research. Finally, the methodology used in this study lends itself to an exploration of genes and metabolites that correlate with or predict a therapeutic response to Combo treatment, but this information is not yet available. As clinical trials progress and biological samples and patient response data become available, we will examine the relationship between gene expression and metabolism in patient-derived samples and clinical effects.

In summary, this study is the first to report the transcriptomic and metabolomic effects of Turso-PB combo in cells from healthy controls and sALS patients. We found that Combo alters the expression of genes involved in ALS-relevant pathways, including mitochondrial function, UPR, nucleocytoplasmic transport, and immune activation. We propose that the modulation of these pathways could underlie neuroprotective effects of Combo in ALS.

## Acknowledgments

We thank Amylyx Pharmaceuticals for providing the compounds used for this study. We thank Dr. Hiroshi Mitsumoto (Columbia University) and the COSMOS initiative for providing the fibroblast and plasma samples utilized in this work. We acknowledge the contribution of the Weill Cornell Genomics Resource Core Facility, and the Medicine Meyer Cancer Center Proteomics & Metabolomics Core Facility. This work was supported by funds from NIH/NINDS grant R35 NS122209 (to GM) and R21 NS104520 (To HK and GM).

## Supplementary Materials

**Figure S1.**
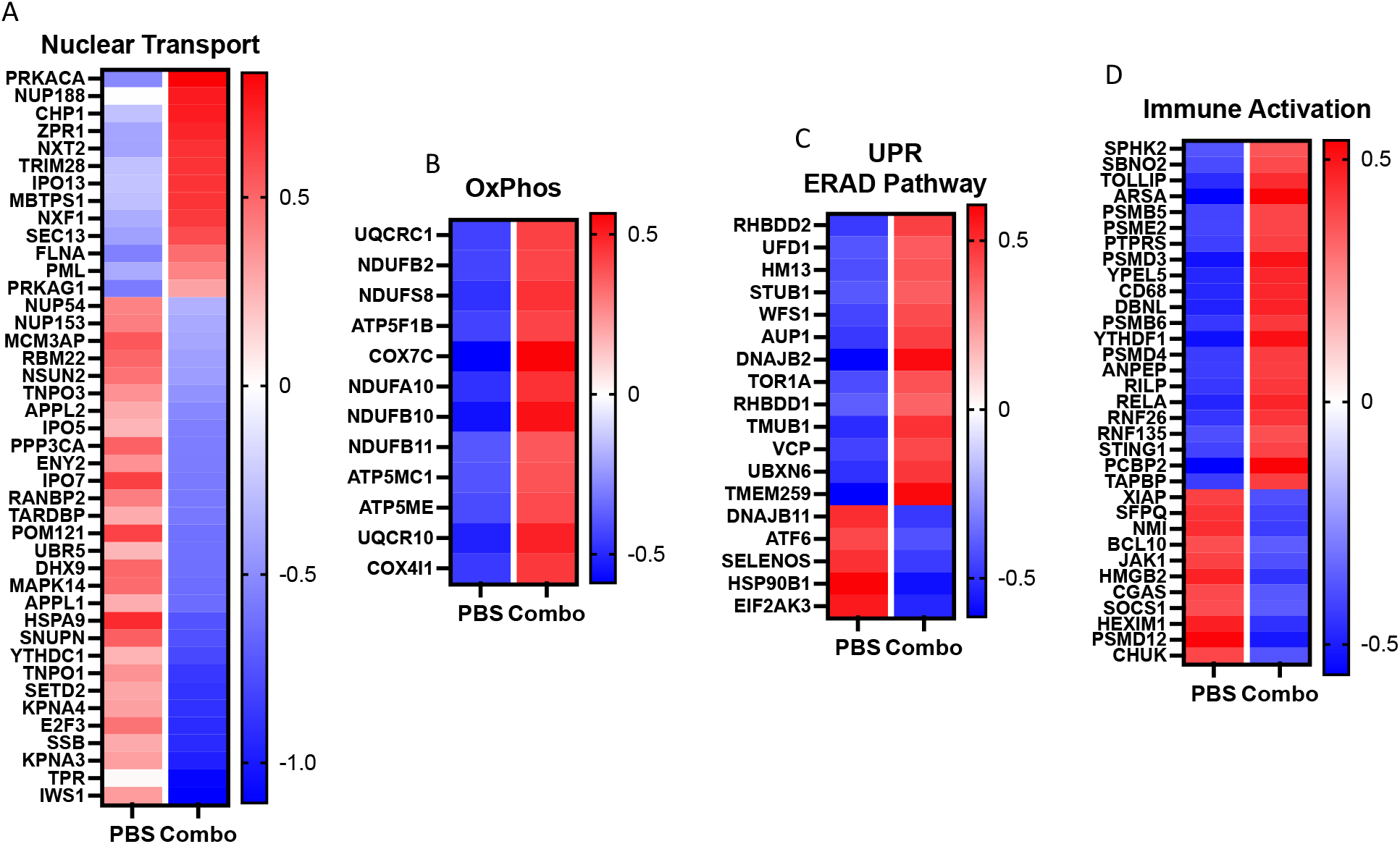
Alterations by Combo treatment in the expression of genes in ALS-relevant pathways. A-D) Heatmaps of Z-scores of DEGs changed by Combo treatment in the nucleo-cytoplastmic transport (A), oxidative phosphorylation (B), unfolded protein response (C), and innate immune activation pathways (D).

**Figure S2.**
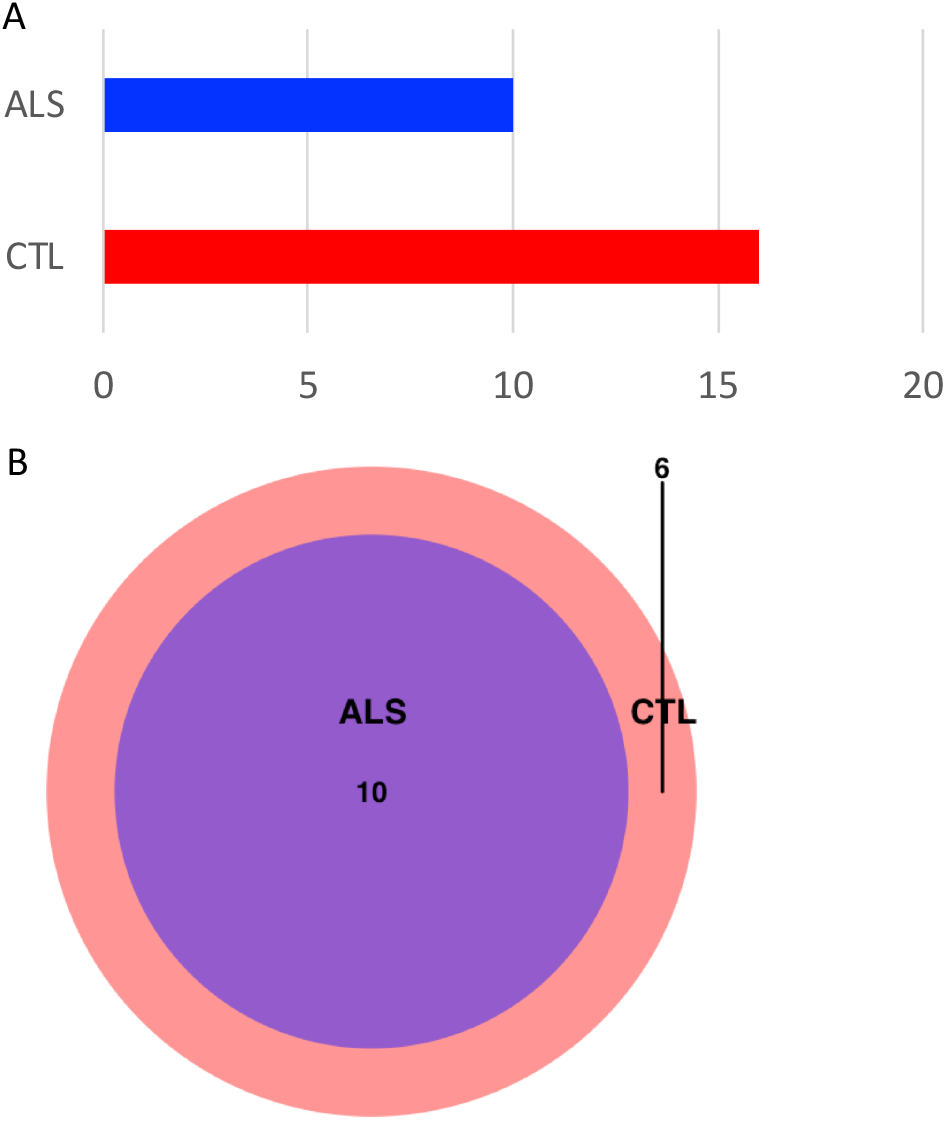
Combo does not have different metabolic effects in sALS and CTL cells. A) Bar graph showing number of significantly different metabolites changed by Combo treatment in ALS and CTL lines (p-value < 0.05). B) Venn diagram of metabolites significantly changed by Combo in ALS and CTL.

**Figure S3.**
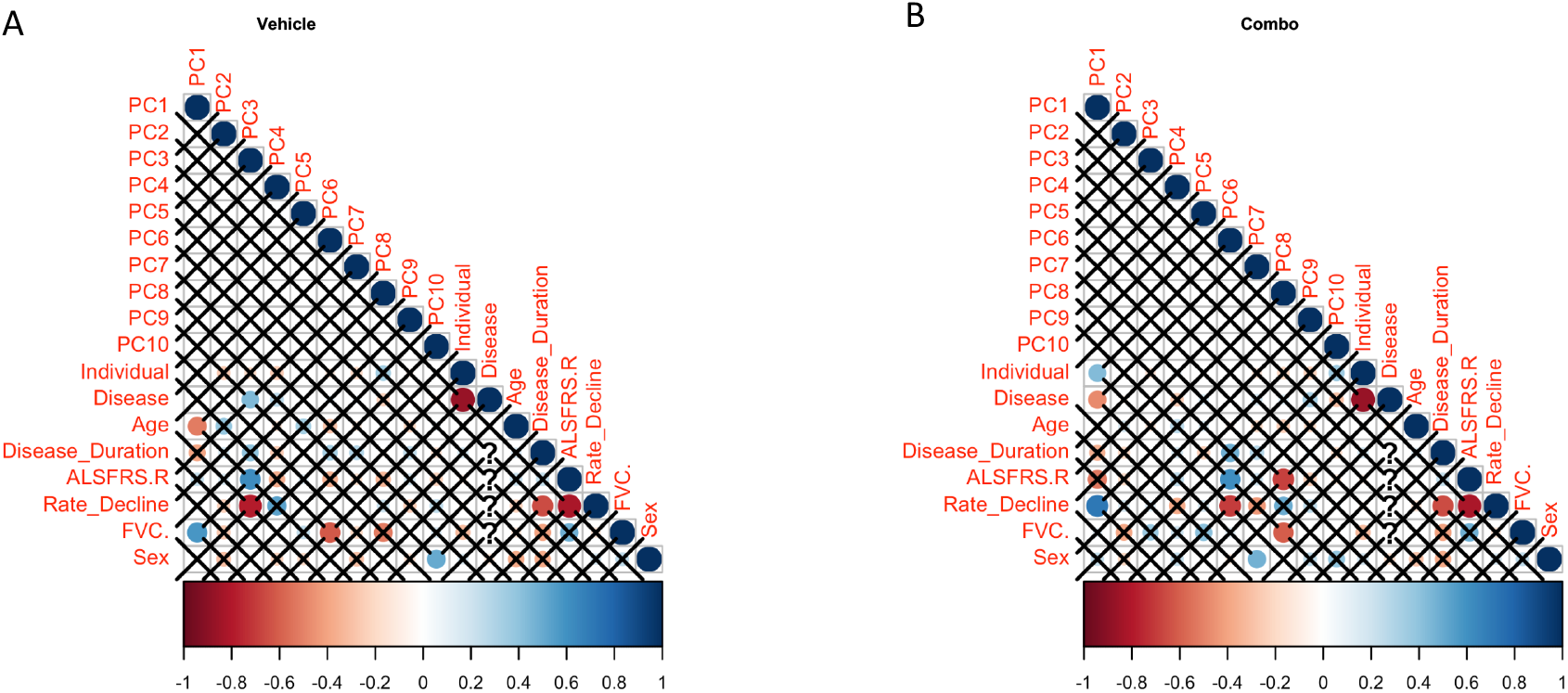
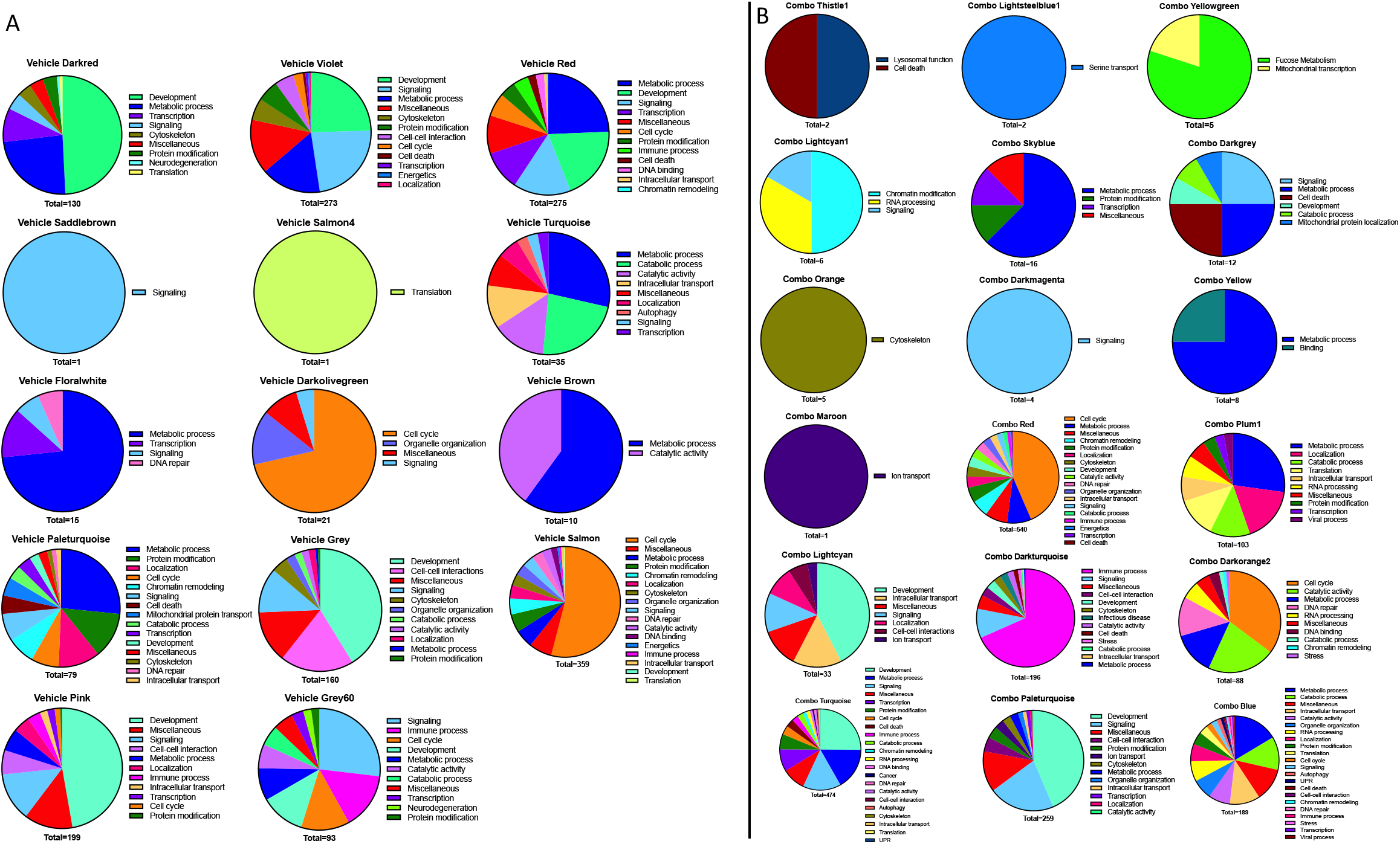
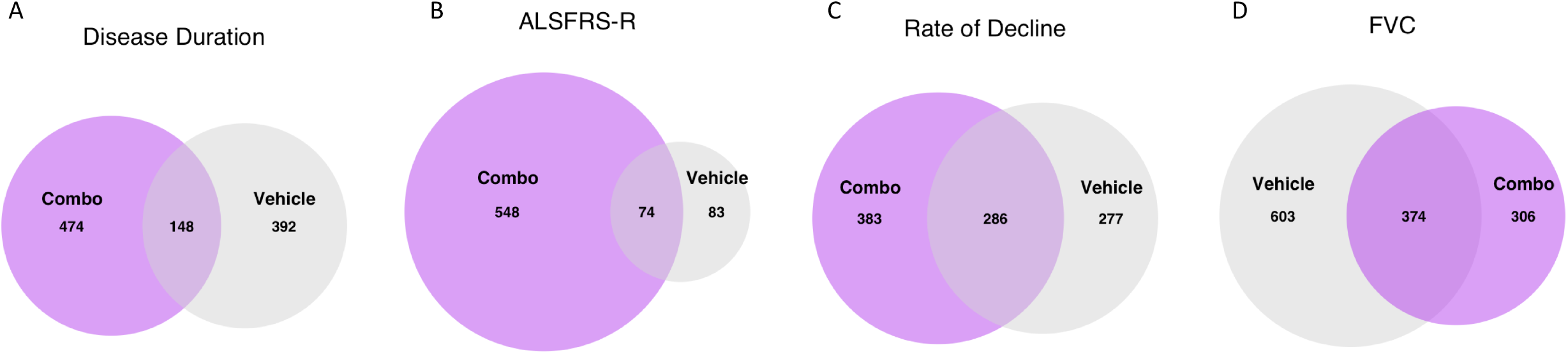
Correlation between gene expression principal components and clinical traits in vehicle and Combo treated cells. A-B) Plots showing correlation coefficients between clinical traits and the first 10 principal components derived from gene expression for the vehicle (A) and Combo (B) networks. Size corresponds inversely to p-value, with X’s denoting correlations that are not statistically significant (adjusted p-value > 0.05), and color corresponds to Pearson’s correlation coefficient.

**Figure S4.**
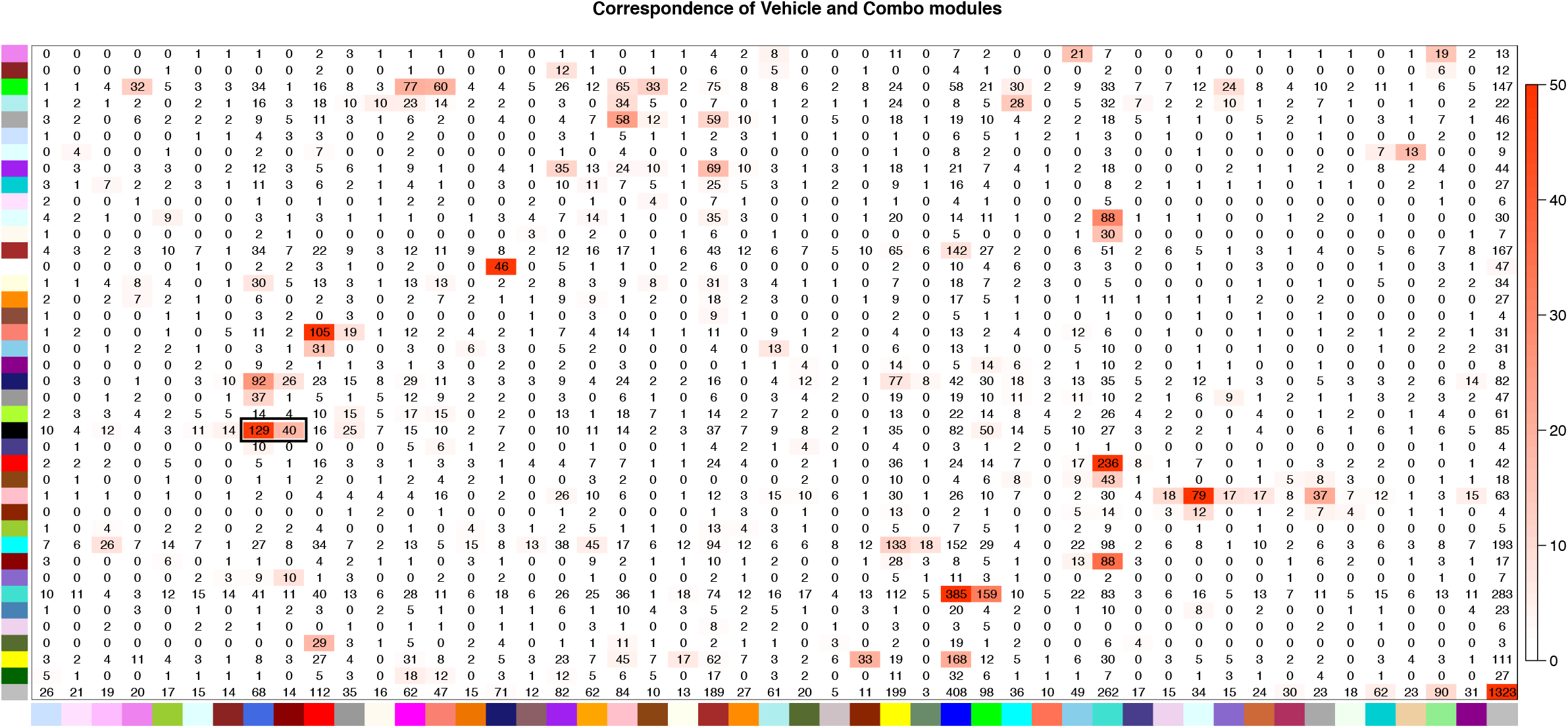
GO pathways associated with clinical traits in vehicle and Combo networks. A-B) Category analysis of all significant GO terms enriched in all significantly trait-associated modules in the vehicle (A) and Combo (B) networks.

**Figure S5**. *GO terms associated with ALS disease traits change after Combo treatment*. A-D) Venn diagrams showing the overlap between all modules significantly associated with clinical traits from the vehicle and Combo networks, for disease duration (A), ALSFRS-R (B), rate of decline (C), and FVC% (D).

**Figure S6**. *Matching of modules in the vehicle and Combo networks*. Table showing correspondence between modules identified in the vehicle (vertical axis) and Combo (horizontal axis) networks. P-values calculated using Fisher’s exact test, color corresponds to −log_10_(p-value). Numbers in each box are the number of overlapping genes found in a pair of modules. The black box identifies the vehicle Black module and its Combo paired modules Darkred and Royalblue.

**Table S1:** Full metabolite identifiers and masses

**Table S2:** Significant differential metabolite abundance results for all treatments compared to PBS

**Table S3:** Differential expression results for all treatments compared to PBS

**Table S4:** GO enrichment results for Combo compared to PBS

**Table S5:** Transcription factor enrichment results for Combo compared to PBS

**Table S6:** Differential expression results for Combo compared to PBS in CTL lines

**Table S7:** Differential expression results for Combo compared to PBS in sALS lines

**Table S8:** GO enrichment results for Combo compared to PBS in CTL lines

**Table S9:** GO enrichment results for Combo compared to PBS in sALS lines

**Table S10:** GO enrichment results for trait-associated modules in vehicle and Combo networks

**Table S11:** Transcription factor enrichment results for the vehicle Black module

